# The human plasma amino acids balance favours HIV replication in primary CD4 T lymphocytes

**DOI:** 10.1101/2024.12.10.627699

**Authors:** Lise Chauveau, Annemarie Fortuin, Arnaud Lecante, Nina Lager-Lachaud, Angéline Tellier, Shravya Honne, Aude Boulay, Fabien P Blanchet, Baek Kim, Floriant Bellvert, Laurence Briant, Jean-Luc Battini

**Affiliations:** RNA viruses and metabolism team, Institut de Recherche en Infectiologie de Montpellier, CNRS UMR 9004, Université de Montpellier, Montpellier, France; Toulouse Biotechnology Institute, Université de Toulouse, CNRS, INRA, INSA, Toulouse, France; MetaToul-MetaboHUB, National Infrastructure of Metabolomics and Fluxomics, Toulouse, France; Center for ViroScience and Cure, Department of Pediatrics, School of Medicine, Emory University, Atlanta, Georgia, USA; Quantitative Biology of Membrane Traffic and Pathogenesis team, Institut de Recherche en Infectiologie de Montpellier, CNRS UMR 9004, Université de Montpellier, Montpellier, France

## Abstract

Cellular metabolism supports all viral replication steps and the metabolic state of infected cells is therefore a key factor influencing viral infections. Human Immunodeficiency virus (HIV) remains latent in resting CD4 T lymphocytes but actively replicates in activated CD4 T cells due to enhanced energy metabolism. Here, using the recently developed Human Plasma-Like Medium (HPLM) that mimics physiological plasma concentration of metabolites, we investigated how this near-physiologic environment modulates HIV-1 infection in primary CD4 T cells. Compared to the conventional culture medium (RPMI), HPLM enhanced HIV-1 infection in CD4 T cells despite similar levels of cell activation, proliferation and expression of viral receptor. In contrast with previous studies in RPMI, HPLM increased infection while decreasing energy metabolism and affecting other non-energetic metabolic pathways. Adjusting levels of several metabolites in RPMI and HPLM, we uncovered that the amino acids balance rather than the energy metabolism favoured HIV-1 replication in this system. Overall, our study used near-physiological conditions to better define metabolic dependencies of viral infections and highlights previously overlooked non-energetic metabolism pathways important for HIV-1 infection.

## Introduction

The Human Immunodeficiency Virus (HIV) is responsible for the worldwide pandemic of Acquired ImmunoDeficiency Syndrome (AIDS). Since the start of the pandemic, 40 years ago, HIV has infected an estimated 88.4 million patients and killed 42.3 million. In 2023, 39.9 million people were still living with HIV^1^. With the development of antiretroviral therapy, AIDS has become a manageable lifelong condition^2^ but no cure or vaccine currently exist^3^. Increasing our understanding of the interaction between HIV-1 and its cellular targets in physiological conditions is crucial to propose better suited treatments.

Cellular metabolism encompasses all pathways involved in the generation or recycling of biomolecules used for energy production, nucleic acid or protein synthesis, regulation of gene expression, redox balance and many other cellular processes. All molecules involved are called metabolites and their homeostasis is tightly regulated. The two main pathways producing energy in the form of ATP are glycolysis and mitochondrial respiration (OXPHOS). HIV-1 mainly infects CD4 T lymphocytes, immune cells crucial for establishing pathogen-specific adaptive immune responses. Upon recognition of pathogen-derived antigens, CD4 T cells are activated and shift from naïve resting cells to actively dividing cells producing large amounts of cytokines. Thus, while naïve T cells keep a low energy supply using low levels of glycolysis and OXPHOS^4^, activated CD4 T cells undergo a profound metabolic reprogramming to meet higher metabolic demands. Increased glucose uptake leads to higher rates of glycolysis to produce energy. Moreover, intermediates in the glycolysis pathway are diverted into other pathways such as the pentose phosphate pathway (PPP) to produce deoxynucleotide triphosphates (dNTPs) and amino acids essential for the cells’ immune functions.

The HIV-1 replication cycle is initiated by the interaction of the viral particle with its receptor (CD4) and co-receptors (CXCR4 or CCR5), leading to the fusion of viral and cellular membranes. Following entry in the cytoplasm, the viral RNA genome is reverse transcribed into a proviral DNA that is imported in the nucleus and integrated in the host genome. From there, viral genes are transcribed and viral proteins are produced which leads to assembly of new viral particles and their budding from the plasma membrane. The different steps of the viral life cycle therefore require the availability of specific metabolites. The increased glycolysis occurring during CD4 T cell activation is associated with higher susceptibility to HIV-1 infection^5–9^. Recent studies even showed that HIV-1 preferentially infects cells with high glycolytic activity, irrespective of their activation status, and blocking glycolysis prevents infection of primary CD4 T cells^7,10^. Both glycolysis and glutaminolysis, which uses glutamine as a carbon source to feed OXPHOS, are essential for HIV-1 replication in these cells^11^.

However, most of these studies were performed *in vitro* using the standard, but not physiologic, RPMI-1640 cell culture medium that was formulated to sustain cell proliferation^12^. Therefore, more physiologically relevant cell culture conditions are needed. Recently, new cell culture media were developed, such as the Human Plasma-Like Medium (HPLM) that closely mimics the metabolite composition of human plasma^13^. Unlike RPMI, HPLM contains 31 polar metabolites absent from commonly used media and carries other components such as glucose, glutamine and amino acids at physiological concentrations. HPLM shapes the immune response in CD4 T cells by influencing the metabolic state of the cells^14,15^.

Here, we investigated how the human plasma metabolic environment affects HIV-1 replication in primary activated CD4 T cells *in vitro*. Compared to RPMI, we found that HPLM potentiated HIV-1 replication by increasing the viral reverse transcription. In contrast with the previous reports performed in RPMI, this increase was associated with lower energy metabolism and lower glycolysis. However, adjusting glucose and glutamine levels in RPMI to match HPLM or inhibiting the energy metabolism using chemical inhibitors did not recapitulate the increased infection observed in HPLM. Instead, adjusting amino acids concentrations in RPMI and HPLM resulted in a similar intermediate infection rate, suggesting that the balance of amino acids in physiological conditions is optimal for HIV-1 replication. Our results therefore highlight the importance of other metabolic pathways that are key for HIV-1 replication in these new near-physiological conditions and were previously overlooked.

## Results

### Human plasma-like metabolic environment potentiates HIV-1 replication in primary activated CD4 T cells

We first wondered if the human plasma-like metabolic environment would affect HIV-1 replication in primary CD4 T cells. CD4 T cells activated in RPMI or in HPLM showed a similar activation by flow cytometry monitoring the activation marker CD25 and cell divisions using Cell Trace (Fig 1A and B). Activated cells were then infected with two doses of HIV-1 NL4-3 virus and virus replication was monitored by flow cytometry looking at expression of the viral Gag protein 4 days post-infection (dpi). Cells cultured in HPLM showed a significantly higher HIV-1 replication compared to those cultured in RPMI with a 2-to 5-fold increase depending on the viral dose used (Fig 1C and D). Treatment with the non-nucleoside reverse transcriptase inhibitor nevirapine (NVP) confirmed that the observed fluorescence correlated with HIV-1 replication (Fig 1C and D). Time course experiments further showed that this increase was not merely due to differences in kinetics of viral replication but to an increased proportion of cells getting actively infected with infection rates reaching up to 65% in HPLM (Fig 1E).

**Figure 1:**
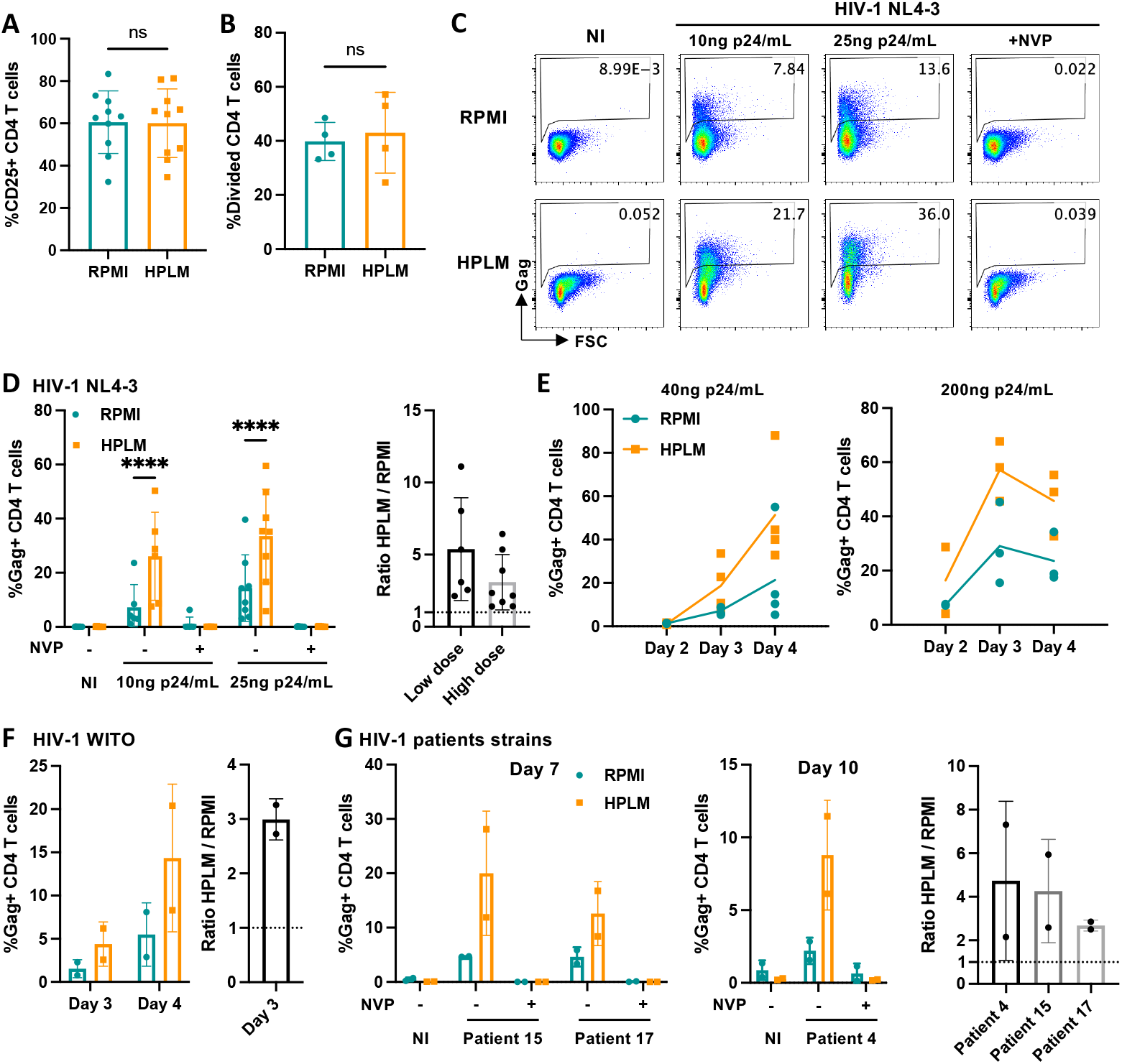
Human plasma-like metabolic environment potentiates HIV-1 replication in primary activated CD4 T cells. Primary CD4 T cells activated in RPMI or in HPLM for 4 days were infected with the indicated strains of HIV-1. **A-B**. At day 4 (**A**) or 5 (**B**) post-activation, flow cytometry analysis of the CD25 activation marker (**A**) and cell proliferation (**B**). **C-E**. Cells were infected with the indicated doses of HIV-1 NL4-3 virus in the presence or absence of the RT inhibitor nevirapine (NVP) as a control. Flow cytometry analysis of the HIV-1 Gag expression at 4 days post-infection (dpi) (**C**, representative donor; **D**, pool) and over time (**E**). NI: non-infected. Raw data (**D**, left) and fold changes in HPLM vs RPMI (**D**, right). **F-G**. Cells were infected with 900ng p24/mL of the transmitted-founder HIV-1 WITO (**F**) or primary isolates from HIV-1 infected patients using the highest possible doses: patient 15: 3,3ng p24/mL, patient 17: 23,5ng p24/mL, patient 4: 2,9ng p24/mL (**G**). Flow cytometry analysis of the HIV-1 Gag expression at the indicated dpi. Raw data (left) and fold changes in HPLM vs RPMI (right). Data show mean±SD of n=10 (**A**), 3 or 4 (**B, E**), at least 6 (**D**) or 2 (**F, G**) donors. Statistical analyses: (**A-B**) Mann-Whitney test; (**D**) 2-way ANOVA followed by Sidak’s multiple comparison test. ns: non-significant; ****: p<0.0001

To confirm the physiological relevance of our results, we next used the transmitted-founder virus HIV-1 WITO as well as 3 primary viral isolates from treatment-naïve patients with high viral loads (patients 4, 15 and 17)^16^. We infected CD4 T cells activated in RPMI or HPLM with these viruses and observed a similar increase in replication (Fig 1F and G) with a 3-fold increase in HIV-1 WITO infection and fold changes ranging from 2 to 7 for the primary isolates revealing variability between donors of the cells used for the infection.

Therefore, HPLM potentiates the replication of several HIV-1 strains including primary isolates in primary activated CD4 T cells.

### HPLM enhances HIV-1 reverse transcription

To investigate the viral replication step affected by the medium, we successively assessed several steps of the viral life cycle using the well-characterised HIV-1 NL4-3 strain. First, we observed that cell surface expression of the HIV-1 receptor (CD4) and co-receptors (CXCR4 and CCR5) were similar when CD4 T cells were activated in each medium (Fig 2A). We then evaluated viral binding by incubating activated CD4 T cells with HIV-1 NL4-3 at 4°C to only allow binding of the HIV-1 viral particle at the cell surface. Quantification of bound particles by HIV-1 Gag p24 ELISA showed no difference between the two media (Fig 2B). Next, we tested if virus fusion at the plasma membrane was affected using the Vpr-Blam assay^17^. Briefly, Vpr, an HIV-1 protein incorporated into the viral particles, is fused with the beta-lactamase (Vpr-Blam). Cells are pre-loaded with CCF2-AM, a cell-permeable fluorescent substrate of beta-lactamase, and infected with viral particles containing Vpr-blam. Upon virus fusion, Vpr-Blam is delivered to the cytosol where it cleaves CCF2-AM, triggering a fluorescent switch quantified by flow cytometry. Using two different doses of HIV-1 NL4-3 Vpr-Blam, we did not observe any difference in fusion between the two media (Fig 2C). We next evaluated reverse transcription (RT) of HIV-1 RNA into DNA, which can be quantified by qPCR using primers specific for the amplification of early and late RT products. CD4 T cells were infected for 18h in the presence or not of the RT inhibitor NVP. We quantified RT products by qPCR on normalised amounts of cellular DNA and calculated copy numbers of early and late RT products using a plasmid standard. We observed a significant 4-fold increase in both early and late RT products in cells cultured in HPLM, suggesting that RT activity is increased in HPLM conditions (Fig 2D).

**Figure 2:**
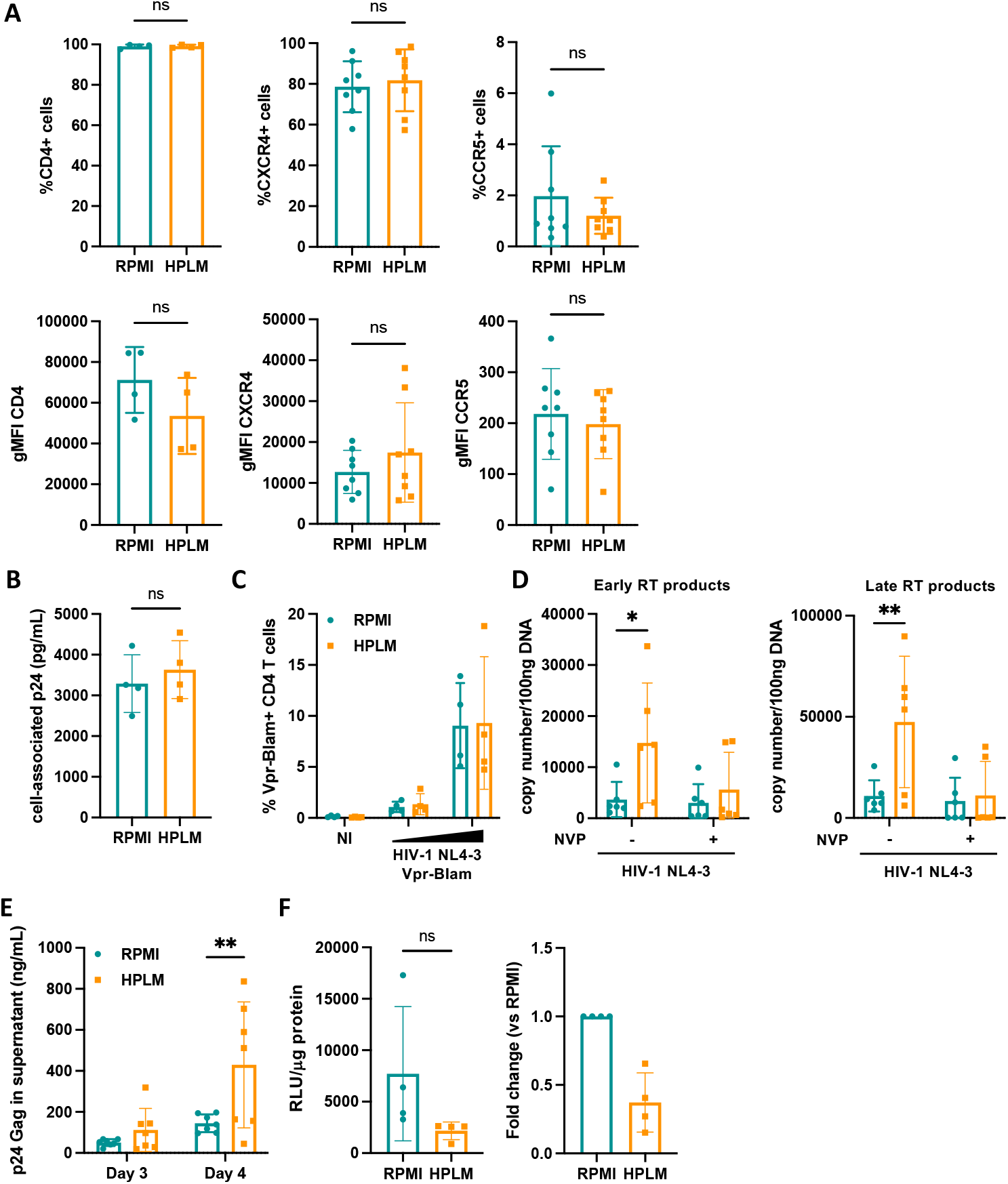
HPLM enhances HIV-1 reverse transcription. Primary CD4 T cells were activated in HPLM or RPMI and infected with the indicated viruses. **A**. Flow cytometry analysis of the HIV-1 receptor CD4 and co-receptors CXCR4 and CCR5 at day 4 post-activation. Percentage of positive cells (upper panels) and geometric mean fluorescence intensity (gMFI) (lower panels). **B**. Cells were placed at 4°C and incubated with 200ng p24/mL HIV-1 NL4-3 for 30min. After extensive washes, the cell-associated virus was quantified by HIV-1 p24 Gag ELISA. **C**. Cells were infected for 2h with 2 doses of HIV-1 NL4-3 Vpr-Blam virus. Flow cytometry analysis of cells where viral fusion occurred using the Vpr-Blam assay. NI: non-infected. **D**. Cells were infected for 2h with 200ng p24/mL DNase treated HIV-1 NL4-3 in the presence or absence of NVP (RT inhibitor) and incubated for 18h before DNA extraction. qPCR for the early and late RT products was performed on normalised quantities of cellular DNA. **E-F**. Cells were infected with 40ng p24/mL HIV-1 NL4-3 for 2h, extensively washed and further incubated. At 3 and 4 dpi, supernatants were collected. The amount of released virus particles was quantified using anti-p24 ELISA (**E**) and their infectivity at 4 dpi was assessed by infecting TZM-bl reporter cells with 5ng p24 of each supernatant (**F**). Raw data (**F**, left) and fold changes in HPLM vs RPMI (**F**, right). Data show mean±SD of n=at least 4 donors. Statistical analyses: Wilcoxon tests (**A, B, F**) or 2-way ANOVA followed by Sidak’s multiple comparison test (**D, E**). ns: non-significant. *: p<0.05; **: p<0.01.

To assess HPLM effects on the late phases of the HIV-1 life cycle, we quantified virus particle released in the supernatant of infected cells by HIV-1 Gag p24 ELISA. Concentration of virus particles were significantly increased in the supernatant of cells infected in HPLM at 4 dpi, correlating with the increased percentage of infected cells in this condition (Fig 2E). However, the infectivity of the virus particles produced assessed on the TZM-bl reporter cell line showed a non-significant decrease (Fig 2F) suggesting that the increased virus particle production could compensate a decrease in particle infectivity.

Therefore, HPLM potentiates HIV-1 replication in activated CD4 T cells primarily by increasing the RT step of the viral life cycle.

### HPLM affects cellular metabolism of primary activated CD4 T cells

HPLM and RPMI formulations differ in the concentrations of three groups of metabolites: carbon sources essential for energy production (glucose, 5mM vs 11mM in RPMI; glutamine, 0.55mM vs 2mM in RPMI); amino acids; and other polar metabolites including some involved in the TCA cycle used in energy metabolism^13^. To investigate how these differences affect CD4 T cells metabolism, we analysed energy metabolism in cells activated in HPLM or RPMI using the ATP rate assay with the Seahorse technology. This method measures the oxygen consumption linked to OXPHOS, and the medium acidification due to lactate export following glycolysis in live cells. It then calculates the overall ATP production rate and distinguishes ATP produced by glycolysis (glycoATP) and by OXPHOS (mitoATP). Activated CD4 T cells had a lower overall ATP production rate and significantly lower glycolysis in HPLM vs RPMI (Fig 3A). We further analysed the metabolism of CD4 T cells activated in HPLM or RPMI by metabolomics using high-resolution mass spectrometry (HRMS). We monitored central metabolism and amino acids by ion chromatography coupled HRMS (IC-HRMS) and liquid chromatography coupled HRMS (LC-HRMS) respectively. PCA analysis indicated significant metabolic differences between the two conditions (Fig 3B). Volcano plot analysis revealed significantly different metabolites (both fold change >2 and p-value<0.1; Fig 3C). Heatmaps for each group of metabolites confidently identified showed specific metabolites variations (Fig 3D). A few metabolites were significantly more abundant in HPLM-activated cells, including phosphorybosyl-pyrophosphate (PRPP), a key metabolite in the pentose phosphate pathway (PPP), and multiple pentose-5-phosphate metabolites, suggesting that the PPP is upregulated in HPLM-activated cells. Moreover, histidine, threonine, glycine, valine, phenylalanine and tyrosine were also elevated in HPLM. Interestingly, this increase did not correlate with higher concentrations of histidine, threonine, phenylalanine and tyrosine in the HPLM medium. In contrast, 21 metabolites were significantly more abundant in RPMI-activated cells, including (i) glutamine and glucose-6-phosphate, the main form of glucose present in cells, reflecting the higher concentrations of these carbon sources in RPMI; (ii) intermediates of glycolysis; (iii) intermediates of the TCA cycle; (iv) intermediates of the hexosamine pathway; and (v) amino acids, mostly reflecting differences in the culture media. We found above that HPLM increases the HIV-1 RT that heavily relies on dNTPs. Specific quantification of cellular dNTPs revealed a decrease in dATP and dTTP and no changes in dCTP or dGTP in HPLM-activated cells (Fig 3E), suggesting that the levels of dNTPs are still sufficient to support RT and are not involved in the phenotype observed.

**Figure 3:**
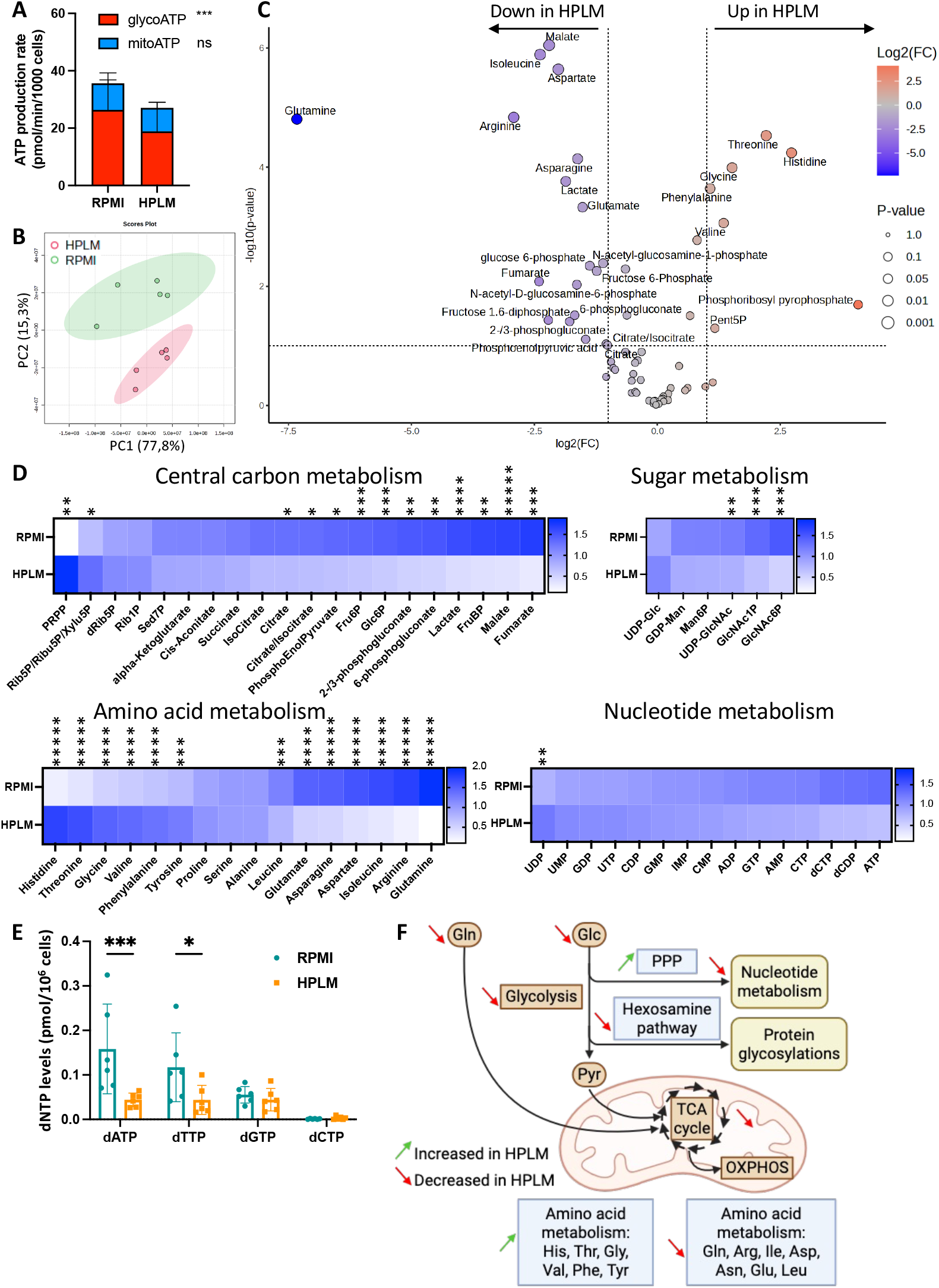
HPLM affects cellular metabolism of primary activated CD4 T cells. General metabolism of primary CD4 T cells activated in HPLM or RPMI was assessed. **A**. Seahorse analysis showing ATP production rate in live cells, including individual contribution of glycolysis (GlycoATP) and OXPHOS (mitoATP). **B-D**. Metabolomics analysis of activated cells from 5 donors by mass spectrometry. PCA analysis comparing the two conditions (**B**). Volcano plot highlighting metabolites with both fold change >2 and p-value <0.1 (**C**). Heatmaps showing changes for each identified metabolite separated into the main metabolic pathways (**D**). **E**. Specific dNTPs quantification in activated cells. **F**. Schematic representation of the metabolic pathways affected by HPLM (Created in BioRender). Data show mean±SD of n=6 (**A, E**) donors. Statistical analysis: 2-way ANOVA followed by Sidak’s multiple comparisons test (**A, E**), or using MetaboAnalyst (**B-D**). ns: non-significant; *: p<0.1; **: p<0.5; ***: p<0.01; ****: p<0.001; *****: p<0.0001.

Overall, these results, summarised in Fig 3F, show that all energy metabolism pathways are down-regulated in cells activated in HPLM, while other metabolic processes such as amino acids and nucleotides metabolisms, the PPP and the hexosamine pathway are altered.

### HPLM enhances HIV-1 infection by providing an optimal amino acid balance

Previous studies have demonstrated that glycolysis and glutaminolysis are crucial for HIV-1 replication in CD4 T cells activated in RPMI^10,11^. However, HPLM has lower concentrations of glucose and glutamine leading to lower energy metabolism but, paradoxically, HIV-1 replicates more efficiently. We hypothesised that lower but physiological concentrations of glucose and glutamine in HPLM could favour HIV-1 replication. We used RPMI without glucose and glutamine supplemented with 5mM glucose and 0.55mM glutamine to match the physiologic concentrations in HPLM (RPMI+Carb). Seahorse analysis showed that cells activated in RPMI+Carb had an intermediate metabolic phenotype that was not significantly different from either RPMI or HPLM (Fig 4A). However, HIV-1 replication in cells activated in RPMI+Carb was similar to RPMI and significantly lower than in HPLM at the two viral doses tested (Fig 4B). Thus, the different energy metabolism in HPLM-activated cells is not linked to the enhanced HIV-1 replication. To confirm that HIV-1 replication still relies on energy metabolism in HPLM, we used three chemical inhibitors targeting different steps in these pathways (summarised in Fig 4C): 2-deoxyglucose (2-DG) targets hexokinase the first rate-limiting enzyme of glycolysis; sodium fluoroacetate (NaFlAc) targets Aconitase and inhibits the TCA cycle shortly after entry of the products of glycolysis; and dimethyl-malonate (DMM) inhibits succinate dehydrogenase, also known as complex II of the mitochondrial respiration chain, the key enzyme to use TCA metabolites in OXPHOS. CD4 T cells activated in RPMI or HPLM were infected with HIV-1 NL4-3 in the presence of these inhibitors. Regardless of the culture medium used, HIV-1 replication was blocked by treatment with 2-DG with an even more pronounced effect in HPLM than in RPMI (Fig 4D). The TCA inhibitors had no significant effect on viral replication with only DMM inducing a non-significant decrease in HIV-1 replication in RPMI.

**Figure 4:**
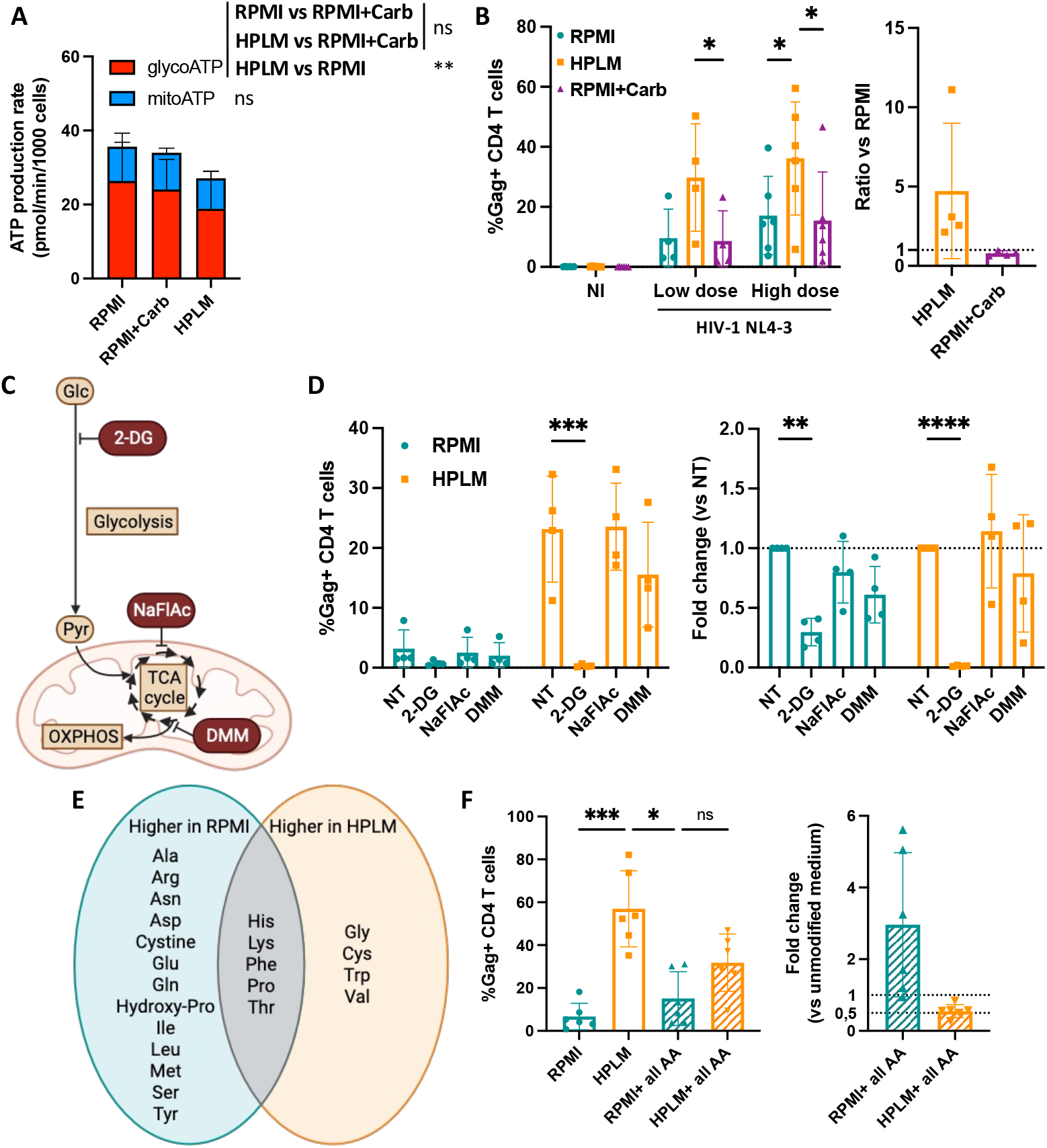
HPLM enhances HIV-1 infection by providing an optimal amino acid balance. **A-B**. Primary CD4 T cells were activated in various media: HPLM, RPMI or RPMI with adjusted concentrations of glucose and glutamine to match the HPLM (RPMI+Carb). **A**. Seahorse analysis showing ATP production rate in live cells, including individual contribution of glycolysis (GlycoATP) and OXPHOS (mitoATP). **B**. Cells were infected as in Fig 1C. Flow cytometry analysis of the HIV-1 Gag expression at 4 dpi. NI: non-infected. Raw data (left) and fold changes in HPLM or RPMI+Carb vs RPMI (right). **C-D**. Cells activated in RPMI or HPLM were infected with 25ng p24/mL HIV-1 NL4-3 in the presence of metabolic inhibitors. **C**. Schematic representation of the energy metabolic pathways targeted by the inhibitors used (Created in BioRender). **D**. Flow cytometry analysis of the HIV-1 Gag expression at 4 dpi. Raw data (left) and fold change compared to the non-treated condition (NT) (right). **E-F**. Cells were activated in various media: RPMI, HPLM, RPMI with adjusted concentrations of amino acids (RPMI+all AA) or HPLM with adjusted concentrations of amino acids (HPLM+all AA). **E**. Schematic representation of the differences in levels of amino acids in the RPMI and HPLM media. Amino acids higher in RPMI were adjusted in HPLM and vice versa. **F**. Flow cytometry analysis of the HIV-1 Gag expression at 4 dpi with 40ng p24/mL. Raw data (left) and fold change of adjusted media with corresponding unmodified medium (right). Data show mean±SD of n= at least 4 (**A, B, D**) or 6 (**E-F**) donors. Statistical analysis: 2-way ANOVA followed by Tukey’s multiple comparisons test (**B**) or Dunnett’s multiple comparisons test (**D**); Friedman test followed by Dunn’s multiple comparisons test (**F**). ns: non-significant; *: p<0.05; **: p<0.01; ***: p<0.001; ****: p<0.0001.

Our metabolomics analysis also revealed a significant difference in the amino acid balance in cells activated in HPLM vs RPMI. We compared the concentrations of all amino acids in the HPLM and RPMI media and identified 4 amino acids with higher concentrations in HPLM and 13 amino acids higher in RPMI (Fig 4E). We therefore adjusted the concentrations of each group of amino acids by adding the 4 in RPMI (RPMI+all AA) and adding the 13 in HPLM (HPLM+all AA). CD4 T cells activated and infected in these media showed an intermediate infection rate that was no longer significantly different between RPMI+all AA and HPLM+all AA (Fig 4F). Fold changes compared to their respective unmodified media revealed that adjusting the amino acids concentrations in RPMI resulted in a 1-to-6-fold increase in HIV-1 replication depending on the donors, while adjusting the amino acids concentrations in HPLM resulted in a 2-fold decrease in HIV-1 replication.

Therefore, while the low glycolysis observed in CD4 T cells activated in HPLM is still crucial for HIV-1 infection, it is not the differences in energy metabolism between RPMI and HPLM but rather the amino acids balance that is involved in the increased HIV-1 replication in cells cultured in HPLM.

## Discussion

In this study, we interrogated the impact of the human plasma metabolic environment on HIV-1 replication in primary activated CD4 T cells. Taking advantage of the recently formulated HPLM as a near-physiologic medium and comparing it to the standard RPMI medium, we show that CD4 T cells activated in HPLM are significantly more susceptible to infection by several strains of HIV-1, including primary isolates from patients. In particular, HPLM increased the reverse transcription (RT) step of the viral replication cycle. Surprisingly, and in contrast to previous studies performed in RPMI, the enhanced HIV-1 replication was associated with a lower energy metabolism, and in particular a lower glycolysis. Several metabolic pathways were affected by the use of HPLM including the Pentose Phosphate Pathway (PPP), the hexosamine pathway and intracellular concentrations of dNTPs and amino acids. The low glycolysis in HPLM-cultured cells was still crucial for HIV-1 infection. However, we show that the optimal amino acids balance in HPLM, rather than the energy metabolism, is responsible for the increased HIV-1 replication in these near-physiological conditions.

HPLM potentiated the replication of several HIV-1 strains including primary isolates from treatment-naïve patients extending our findings to the most physiologically-relevant replication-competent viruses. This increased replication was observed in cells showing similar cellular activation or proliferation. The kinetics of replication in our assays were also similar between the two media with the peak of infection reaching up to 65% in HPLM vs 45% in RPMI suggesting that it is the number of cells susceptible to infection that was increased in HPLM. These results are concordant with previous findings showing that CD4 T cells proliferation^7^ or activation^10^ status do not predict susceptibility to HIV-1 infection. Instead, the former study found that the metabolic state of the cells, and in particular their uptake of glucose, is important for HIV-1 RT. Similarly, we observed a 3-fold increase in HIV-1 RT at 18 hours post-infection in HPLM-activated CD4 T cells, which likely contributed to the increased viral replication observed at 4 days post-infection. The RT step of the viral life cycle is highly dependent on cellular metabolism which conditions availability of the necessary dNTPs^18^. Our metabolomics analysis revealed an increased PPP in HPLM-cultured cells and this pathway provides precursors for *de novo* dNTPs synthesis. However, our intracellular dNTPs quantification unexpectedly revealed decreased dNTPs concentrations in HPLM-cultured cells, particularly dTTP and dATP. HPLM was previously shown to inhibit *de novo* pyrimidine synthesis in cancer cell lines by providing urea, a specific inhibitor of this pathway^13^. In primary CD4 T cells, HPLM could therefore both increase the PPP and block dTTP and dATP synthesis via unknown mechanisms worth exploring in the future. Our results show that this lower concentration of dNTPs is still sufficient to support HIV-1 RT in HPLM-activated CD4 T cells and suggest that other metabolites modulated by HPLM are important for HIV-1 RT. Moreover, whether this result reflects more RT per infected cell or an increase in the number of cells where the RT step is efficient could not be determined in our assays. We can also not exclude that another unexamined step of the viral replication cycle, such as integration of the viral genome or cell-to-cell viral transmission, may contribute to the increased HIV-1 replication in HPLM conditions.

HPLM decreases the hexosamine pathway while increasing the PPP in activated CD4 T cells. The hexosamine pathway produces UDP-N-acetyl-D-glucosamine (UDP-GlcNAc), the substrate for both N-and O-protein glycosylations, and is therefore important for this protein post-translational modification. In the context of HIV-1 infection, glycosylation of the Sp1 transcription factor was shown to inhibit HIV-1 transcription from the LTR both in T cell lines and in primary CD4 T cells^19^. In HPLM, the decreased hexosamine pathway could therefore decrease Sp1 glycosylation and increase HIV-1 transcription. The HIV-1 envelope gp120 is also glycosylated which increases viral particle infectivity while decreasing recognition by neutralising antibodies^20^. In HPLM, this could explain the trend of decreased viral particles infectivity observed in Fig 2F which warrants further investigations. As previously mentioned, the PPP provides precursors for nucleotides synthesis but also precursors for amino acids synthesis, and NADPH for oxidative stress management. While we show that nucleotide levels are unaffected or decreased in HPLM, other downstream effects of the increased PPP in HPLM could be involved in the enhanced HIV-1 replication. Interestingly, the metabolic phenotype we observe in HPLM – downregulated glycolysis with increased PPP activity - is a hallmark of the establishment of HIV-1 latency^21^. HPLM could therefore serve as a valuable model for studying HIV-1 latency, the current main hurdle in the search for an HIV cure, and exploring therapeutic strategies such as “shock and kill”.

Glycolysis and OXPHOS are key pathways previously shown to support HIV-1 infection in CD4 T cells activated in RPMI ^10,11,22^. Inhibitors of these pathways were reported to significantly reduce HIV-1 infection^10^. However, despite exhibiting lower energy metabolism, CD4 T cells activated in HPLM showed a higher susceptibility to HIV-1 infection. Our findings reveal that decreased energy metabolism is not directly responsible for the increased infection, as adjusting glucose and glutamine levels in RPMI to match HPLM did increase HIV-1 replication to levels found in HPLM-activated CD4 T cells. In line with previous reports, we show that the glycolysis inhibitor 2-DG decreased HIV-1 replication, with an even more pronounced effect in HPLM than in RPMI. Our results therefore extend these previous findings to CD4 T cells cultured in HPLM where the limited glucose availability might increase 2-DG efficacy. In contrast, we did not observe a significant effect with TCA cycle or OXPHOS inhibitors such as DMM that lead to a non-significant decrease in HIV-1 replication in RPMI and only a modest effect in HPLM. This result contradicts recent studies^22^ and suggests that HIV-1’s reliance on OXPHOS might differ between the two media.

Amino acids are another group of metabolites that are present at different concentrations in HPLM and in RPMI. Our metabolomics analysis revealed that amino acids metabolism was differentially regulated in cells activated in each medium suggesting that they could be of importance for HIV-1 replication. When adjusting the concentrations of amino acids, adding some in RPMI and others in HPLM, we observed an intermediate infection rate that was similar between the two adjusted medium. Therefore, amino acids that we added into RPMI could have pro-viral effects and amino acids added into HPLM could have anti-viral effects. Some of the amino acids in both lists already have pro- or antiviral effects described in the context of HIV-1 infection. For instance, tryptophane present in HPLM enhances HIV-1 transcription in cell lines and viral reactivation in latently infected cells derived from people living with HIV^23^. Moreover, arginine present in RPMI can be used to produce polyamines that have an antiviral role by blocking HIV-1 fusion^24^. The effects of individual amino acids and their combinations on HIV-1 replication in the context of HPLM requires further investigation.

In conclusion, our study identifies the amino acid balance in human plasma as a crucial factor favouring HIV-1 replication in primary CD4 T cells. It further highlights the critical role of metabolic pathways other than glycolysis and OXPHOS that have been overlooked in this context and will deserve attention in the future to identify new therapeutic targets. We also argue for the importance of using physiologically relevant models, which encompasses primary cells and more physiologic culture media, to investigate host-pathogen interactions, especially when looking at metabolic dependencies. This needs to be particularly considered for antiviral drugs development and for the study of latent HIV-1-infected CD4 T cells which account for HIV-1 persistence and incurable life-long infection in people living with HIV.

## Materials and methods

### Cells

Cells were kept in culture in the following media: classical RPMI-1640 medium (Gibco); newly developed Human Plasma-Like Medium (HPLM, Gibco); RPMI without glucose and glutamine (PAN Biotech) where glucose was added at 5mM and glutamine at 0.55 mM to match the concentrations in HPLM (RPMI+Carb); RPMI (Gibco) complemented with various amino acids to match the concentrations in HPLM (RPMI+all AA); or HPLM (Gibco) complemented with various amino acids to match the concentrations in RPMI (HPLM+all AA). RPMI, RPMI+Carb and RPMI+all AA were supplemented with 10% heat-inactivated fetal calf serum (FCS) and 1% PenStrep (Gibco). HPLM and HPLM+all AA were supplemented with 10% heat-inactivated dialysed FCS (Gibco) and 1% PenStrep (Gibco).

PBMCs were isolated from the blood of healthy donors by Ficoll centrifugation. The blood was provided by the EFS (Etablissement Francais du Sang, the official French blood bank). CD4+ T lymphocytes were isolated from PBMCs by positive selection using magnetic beads (EasySep Human CD4 positive selection kit II, StemCell Technologies) and activated using ImmunoCult human CD3/CD28 T cell activator cocktail (StemCell Technologies) then grown in the indicated medium in presence of 100U/mL IL-2 (StemCell Technologies) for 4 days before being used in the various experiments. In all experiments, IL-2 was added in the culture medium at a final concentration of 100U/mL to support cell survival and growth. HEK293T and TZM-bl cells were cultured in DMEM supplemented with 10% FCS and 1% PenStrep (Gibco).

### Reagents

The reverse transcription inhibitor nevirapine (NVP) was from the NIH AIDS reagents program. The following metabolic inhibitors were used: 2-deoxy-glucose (2-DG, 5mM; Sigma); dimethyl malonate (DMM, 10mM; Sigma); sodium fluoroacetate (NaFlAc, 10μM; CliniSciences).

### Viruses

HIV-1 NL4-3 and WITO were generated by transfection of pNL4-3 and pWITO.c/2474, respectively, in HEK293T cells. Briefly, HEK293T cells were seeded in T175 flasks and transfected the next day using PEIpro (Polyplus). After overnight incubation, the medium was changed and the supernatant was collected 24h, 32h and 48h later. The three harvests were pooled and concentrated by ultracentrifugation through a 20% sucrose cushion at 100,000g for 2h then resuspended in PBS. Primary isolates from treatment-naïve patients were a kind gift from Nathalie Arhel and have been previously described^16^. The concentration of the viral stock was determined using HIV-1 Gag p24 ELISA (Lenti-X™ p24 Rapid Titer Kit #631476, Takara; or Innotest HIV Ag mAb, Fujirebio). The doses used in all experiments were chosen by infecting primary CD4 T cells to reach appropriate levels of infection and were adjusted for each viral production.

### HIV-1 infection

150 000 primary CD4 T cells were seeded in round-bottom 96-well plates and infected in a final volume of 100uL with 10, 25, 40 or 200 ng p24/mL of HIV-1 NL4-3 WT as indicated, or 900ng p24/mL HIV-1 WITO, or viruses from patients at the indicated concentrations. When indicated, the reverse transcriptase inhibitor NVP or the metabolic inhibitors were added in the wells at the time of infection.

For analysis of HIV-1 reverse transcription, the input virus was first treated with RQ1 DNase (Promega) for 1h at 37°C before adding to the cells to remove left-over plasmid from the production transfection. Two hours after infection, the cells were washed in PBS to remove input virus and kept in culture for an additional 18h.

For analysis of the virus particle release in the supernatant, the cells were washed twice in PBS 2h after infection in order to remove the input virus and kept in culture for 3 or 4 days.

### Flow cytometry

To assess activation of the CD4 T cells prior to infection, activated cells were stained with anti-CD25 FITC (Biolegend) in PBS+2%FCS+1mM EDTA (FACS Buffer) for 30min at 4°C and washed twice in FACS Buffer.

To quantify proliferation of the CD4 T cells, cells were stained directly after isolation from PBMCs with the CellTrace Far Red (ThermoFischer Scientific) in PBS for 20min at 37°C. Medium was added to the cells to dilute the dye and further incubated for 5min at RT. Cells were then centrifuged, resuspended in the appropriate volume of each medium and activated with the ImmunoCult activation cocktail. Proliferation was monitored by flow cytometry at day 5 post-activation by looking at loss of fluorescence due to cell division and the percentage of cells having divided at least once was calculated.

To assess the infection rate at the indicated days post-infection, cells were stained with a fixable viability dye (LIVE/DEAD fixable far red viability stain, ThermoFisher Scientific) and fixed in 4% PFA for 15min at room temperature. Intracellular Gag staining was performed by permeabilising cells in FACS Buffer + 0.5% Triton X-100 for 20min at RT then staining with the FITC-conjugated KC-57 anti-Gag antibody (Beckman-Coulter) in FACS Buffer for 30min at 4°C. After two washes in FACS buffer, the infection was analysed by flow cytometry looking at the percentage of cells expressing Gag.

All samples were analysed with a NovoCyte Flow Cytometer (Agilent).

### Binding assay

150 000 primary CD4 T cells were seeded in round-bottom 96-well plates and placed at 4°C for 2h. They were then incubated with 200ng p24/mL HIV-1 NL4-3 for 30min at 4°C. Unbound viral particles were washed 4 times in PBS, cells were lysed and bound viral particles were quantified using HIV-1 Gag p24 ELISA (Innotest HIV Ag mAb, Fujirebio) according to manufacturer’s instructions.

### Fusion assay

Viral fusion was assessed as previously described^17^. Briefly, ultracentrifuged HIV-1 NL4-3 virions containing the Blam-Vpr fusion protein (a kind gift from Caroline Goujon) were used to infect primary CD4 T cells. After 2h, cells were washed in CO_2_-independent medium supplemented with 10% FCS then incubated in CCF2-AM loading solution (CCF2-AM kit, ThermoFisher scientific) containing 2.5mM Probenecid (Sigma) for 2h at room temperature. Cells were then washed in CO_2_-independent medium supplemented with 10% FCS and 2.5mM Probenicid and further incubated overnight in the same medium. The next day, cells were washed with PBS, fixed in 4% PFA for 15min, resuspended in FACS Buffer and analysed with a Novocyte Flow Cytometer looking at CCF2-AM in the AmCyan channel and cleaved CCF2-AM in the Violet channel.

### Reverse transcription assay

Infected cells DNA was extracted using the DNeasy Blood and Tissue kit (Qiagen). Between 50-100ng of total DNA were used with the LightCycler 480 SYBR Green I Master mix (Roche) and amplified on a BioRad CFX Opus 384 Real-time PCR system. Primers for HIV-1 NL4-3 ssDNA early products were: M667 (5′-GGCTAACTAGGGAACCCACTG -3’) and AA55M (5′-GCTAGAGATTTTCCACACTGACTAA -3′) and for strand transfer late products: M667 and M661 (5′-CCTGCGTCGAGAGAGCTCCTCTGG -3′). PCR conditions were as follows: 95°C for 10min then 40 cycles of 95°C 15s, 65°C 15s and 70°C 20s. Quantities of cellular DNA were calculated using a plasmid standard and normalised to the amount of DNA used.

### Virus particle release assay

At days 3 and 4 post-infection, supernatant of infected cells was harvested after cell centrifugation and frozen at -80°C. p24 titration was performed using an HIV-1 Gag p24 ELISA kit according to the manufacturer’s protocol (Lenti-X™ p24 Rapid Titer Kit #631476, Takara; or Innotest HIV Ag mAb, Fujirebio).

### Viral particle infectivity assay

Infectivity of the viral particles found in cell supernatants and quantified as described above was assessed on the TZM-bl reporter cell line that expresses Luciferase under the HIV-1 LTR promoter. 10 000 TZM-bl cells were seeded in flat-bottom 96-well plates. The next day, medium was changed and 5ng p24 of each supernatant was added to the cells in duplicate. 48h post-infection, cells were lysed in Passive Lysis Buffer (Promega) and Firefly Luciferase activity was quantified using Genofax A (Yelen).

### Seahorse ATP rate assay

Seahorse analyses of the energy metabolism in activated CD4 T cells were performed using the ATP rate assay (Seahorse Biosciences, Agilent) following manufacturer’s protocol. Briefly, cells were counted, resuspended in Seahorse RPMI culture medium and seeded in six replicates in XF96 plates pre-coated with 0.1mg/mL poly-D-lysine. Cells were then incubated for a minimum of 1h in a CO_2_-free incubator at 37°C before the plate was loaded in the Seahorse XF96e analyser. The protocol for the ATP rate assay was used following manufacturer’s instructions with the following injections: (1) oligomycin (1μM); and (2) rotenone and antimycin A (1μM each) in ports A and B respectively. The ATP rate assay analysis was performed using the Seahorse Analytics online software.

### Metabolomics analyses by IC- and LC-HRMS

For metabolomics analyses, CD4 T cells were activated in RPMI or HPLM for 5 days then washed in cold PBS and cell pellets were snap-frozen in liquid nitrogen and stored at -80°C until metabolites extraction. Sampling buffer (acetonitrile:methanol:water at a ratio of 4:4:2 + 1.25mM formic acid) was prepared and cooled to -20°C 2h before extraction. Cell pellets were resuspended in 1mL sampling buffer with IDMS controls and incubated for 1h at -20°C. Cell debris were discarded by centrifugation at 2000g for 5min at 4°C. Supernatants containing metabolites were transferred into a new tube, dried using a Speedvac and stored at -80°C until mass spectrometry analyses.

Central metabolites were separated on an ionic chromatography column IonPac AS11 (250 × 2 mm i.d.; Dionex, CA, USA). Mobile phase used was a gradient of KOH at a flow rate of 380 μL/min. Mobile phase was varied as follows: 0 min: 7 mM, 1 min: 7 mM, 9.5 min: 15 mM, 20 min: 15mM, 30 min: 45 mM, 33 min: 70mM, 33.1 min: 100mM and 42 min: 100mM. The column was then equilibrated for 5 min at the initial conditions before the next sample was analyzed. Time of analysis is 50 minutes. The volume of injection was 15μL. High-resolution experiments were performed with an ICS5000+, ion chromatography system (Dionex, CA, USA) system coupled to an Orbitrap Qexactive+ mass spectrometer (Thermo Fisher Scientific, Waltham, MA, USA) equipped with a heated electrospray ionization probe. MS analyses were performed in negative FTMS mode at a resolution of 70 000 (at 400 m/z) in full-scan mode, with the following source parameters: the capillary temperature was 350 °C, the source heater temperature, 350 °C, the sheath gas flow rate, 50 a.u. (arbitrary unit), the auxiliary gas flow rate, 10 a.u., the S-Lens RF level, 65 %, and the source voltage, 2.75 kV. Metabolites were determined by extracting the exact mass with a tolerance of 5 ppm.

For amino acid analysis, amino acids were separated on a PFP column (150 x 2.1 mm i.d., particle size 5 μm; Supelco Bellefonte, PEN, USA). Solvent 1 was 0.1% formic acid in H2O and solvent B was 0.1% formic acid in acetonitrile at a flow rate of 250 mL/min. The gradient was adapted from the method used by Boudah et al. Solvent B was varied as follows: 0min: 2%, 2 min: 2%, 10 min: 5%, 16 min: 35%, 20 min: 100% and 24 min: 100%. The column was then equilibrated for 6 min at the initial conditions before the next sample was analysed. The volume of injection was 5μL. For LTQ-Orbitrap High-resolution experiments were performed with Vanquish HPLC system coupled to an LTQ Orbitrap Velos mass spectrometer (Thermo Fisher Scientific, Waltham, MA, USA) equipped with a heated electrospray ionization probe. MS analyses were performed in positive FTMS mode at a resolution of 60 000 (at 400 m/z) in full-scan mode, with the following source parameters: the capillary temperature was 275 °C, the source heater temperature, 250 °C, the sheath gas flow rate, 45 a.u. (arbitrary unit), the auxiliary gas flow rate, 20 a.u., the S-Lens RF level, 40 %, and the source voltage, 5 kV. Metabolites were determined by extracting the exact mass with a tolerance of 5 ppm.

Combined analysis was performed using MetaboAnalyst v6.0.

### dNTPs quantification

4×10^6^ CD4 T cells activated in RPMI or HPLM for 4 days were washed with cold PBS then lysed in ice cold 65% methanol, vortexed vigorously and incubated at 95°C for 3min. After cooling down the samples on ice for 1 min, they were centrifuged at 14 000rpm for 3 min and the methanol supernatant was collected into a new tube, dried with a Speedvac and stored at -80°C. The extracted dNTP samples were analyzed by the RT-based primer extension dNTP assay^25^. Briefly, the dNTP samples were used for each 20 μl single nucleotide incorporation primer extension reaction by HIV-1 RT. The dNTP levels in each sample were determined within the linear ranges of the dNTP incorporation (2∼32% primer extension), and the primer extension levels in each dNTP sample from a known number of cells were converted into dNTP quantities (pmol) per 1×10^6^ cells.

## Acknowledgments

The authors thank Lucile Espert and Caroline Goujon for sharing reagents and advice, Nathalie Arhel for sharing the HIV-1 patients’ strains, and all team members for technical help and advice. This work was supported by the French AIDS National Research Agency ANRS (ECTZ134139 to J.-L. B.), the U.S. National Institute of Health NIH (R01 AI136581 and R01 AI162633 to B. K.) and used METABOHUB project funded by the program Investments for the Future of the French National Agency for Research (N. L.-L. and F.B.; ANR 11-INBS-0010). L.C. was supported by the French AIDS National Research Agency ANRS (ECTZ134113) then by CNRS, A.L. was supported by the French Sidaction (#12619 and #13564), A.B. was supported by an ANRS grant attributed to F.P.B. (ECTZ192763), A.T. was supported by a French Sidaction grant attributed to L.C. (#20224), L.B. is supported by CNRS and F.P.B. and J.-L.B. by Inserm.

